# Partition complex structure can arise from sliding and bridging of ParB dimers

**DOI:** 10.1101/2022.12.01.518708

**Authors:** Lara Connolley, Lucas Schnabel, Martin Thanbichler, Seán M. Murray

## Abstract

Chromosome segregation is vital for cell replication and in many bacteria is controlled by the ParAB*S* system. A key part of this machinery is the association of ParB proteins to the *parS*-containing centromeric region to form the partition complex. Despite much work, the formation and structure of this nucleoprotein complex has remained unclear. However, it was recently discovered that CTP binding allows ParB dimers to entrap and slide along the DNA, as well as leading to more efficient condensation through ParB-mediated DNA bridging. Here, we use semi-flexible polymer simulations to show how these properties of sliding and bridging can explain partition complex formation. We find that transient ParB bridges can organise the DNA into either a globular state or into hairpins and helical structures, depending on the bridge lifetime. In separate stochastic simulations, we show that ParB sliding reproduces the experimentally measured multi-peaked binding profile of *Caulobacter crescentus*, indicating that bridging and other potential roadblocks are sufficiently short-lived that they do not hinder ParB spreading. Indeed, upon coupling the two simulation frameworks into a unified sliding and bridging model, we find that short-lived ParB bridges do not hinder ParB sliding and the model can reproduce both the ParB binding profile as well as the condensation of the nucleoprotein complex. Overall, our model clarifies the mechanism of partition complex formation and predicts its fine structure. We speculate that the DNA hairpins produced by ParB bridging underlie the secondary function of ParB to load the Structural Maintenance of Chromosome (SMC) complex onto the DNA.

---

**F**aithful chromosome segregation is essential for the efficient replication of cells. For this, bacteria chromosomes and low-copy plasmids employ active partitioning systems, with the ParAB*S* system being one of the most widespread (1–3). This system consists of three components: centromeric-like *parS* sequences and two proteins, ParB which forms dimers that bind specifically to the *parS* sequence, and ParA, an ATPase, the activity of which is stimulated by ParB (4, 5).

ParB spreads to several kilobases of DNA surrounding the *parS* sites, which in bacteria are concentrated close to the origin of replication (6). This spreading is essential in order for these systems to function, though the degree of spreading varies substantially between systems (7–13). In any case, the result is believed to be a nucleoprotein complex, the partition complex, that is clearly visible using fluorescence microscopy. Originally, spreading was proposed to be due to the formation of a nucleoprotein filament extending out from the *parS* site (7, 8, 14). However, it was subsequently shown that there are too few ParB proteins to form such large structures (10). Instead, ParB was found *in vitro* to condense DNA through non-specific DNA binding and the formation of protein bridges (10, 15–19).

These results motivated modelling studies of partition complex formation. In particular, the spreading and bridging model proposed that the partition complex forms through a combination of long-range (3D) bridging and short-range (1D) nearest-neighbour interactions (20). However, this model was subsequently argued to be incompatible with the binding profile of ParB from F plasmid (11). Instead, it was proposed that the observed profile is due to the spatial caging of ParB around the nucleating *parS* site, due to non-specific and transient interactions, and the polymeric nature of the DNA (11, 21, 22).

Recently, it has been demonstrated that ParB exhibits *parS*-dependent CTPase activity that is required for correct partition complex formation and dynamics (13, 23–27). CTP-bound ParB dimers were shown to load onto DNA at *parS* sites and subsequently slide along the DNA strand before eventually dissociating. It was also shown *in vitro* that CTP binding allows ParB bridging to occur at physiological concentrations (much lower than that required in the absence of CTP (10, 15)) and leads to efficient DNA condensation (28, 29). These results fundamentally change our understanding of how ParB can spread and suggest that the previous models need to be reevaluated. ParB sliding may also have additional relevance for chromosomal ParABS systems, which typically have several genomically separated *parS* sites and, as a result, more than one peak in the ParB binding profile (9, 10, 12, 13, 30–33), yet have a single visible partition complex per origin in wild-type cells (10, 34). The effect of multiple *parS* sites on partition complex formation has not yet been studied computationally.

Here, we investigate the role of ParB sliding and bridging in partition complex formation using semi-flexible polymer and reaction-diffusion simulations. Focusing on the ParAB*S* system of *Caulobacter crescentus*, we first show that different ParB bridge lifetimes lead to distinctly different polymer conformations. We then study the short-lifetime regime in which distinct DNA structures (hairpins and helices) form and show how these structures result in the condensation of ParB-coated DNA. We then use stochastic simulations to show that ParB sliding can reproduce the multi-peaked ParB distribution seen experimentally and explore the effects of roadblocks on sliding. Finally, we combine ParB bridging and sliding in coupled polymer/reaction-diffusion simulations and show that bridging does not inhibit ParB sliding for sufficiently short bridge lifetimes. Overall, our work supports a new model of partition complex formation in which ParB dimers load onto the DNA at *parS* sites before sliding diffusively along the DNA. Genomically distant, but spatially proximal, ParB dimers interact to form transient bridges that organise the DNA through the formation of hairpin and helical structures. We speculate that these structures facilitate the additional function of ParB to load Structural Maintenance of Chromosomes (SMC) complexes onto the chromosome.

## Results

### Semi-flexible polymer model of ParB bridging

In order to obtain a realistic model of partition complex formation, we use a semi-flexible lattice polymer model of the centromeric region of *C. crescentus* in which every monomer corresponds to 20 bp, the approximate footprint of a ParB dimer (27, 35) (Fig. 1A). In particular, we use a kinetic implementation of the Bond Fluctuation Method (BFM) (36), an ergodic polymer model that reproduces Rouse polymer dynamics. Since the DNA is stiff at this scale, we introduce an energy cost for bending to obtain the experimentally measured persistence length of *l*_*p*_ ∼120 bp (SI Fig. 1A) (37). The BFM is well suited for this as it allows a large set of bond angles (36) and can therefore implement stiffness more realistically than models that only allow 0° or 90° bond angles. Bridging between DNA-bound ParB dimers is implemented by allowing any two spatially proximal, non-neighbouring monomers of the polymer to form a bridge with a probability dependent on their ParB occupancy. Each dimer/monomer can only bridge one other dimer/monomer at a time (in the following ‘dimer’ will always refer to a ParB dimer and ‘monomer’ to a monomer of the simulated DNA polymer). Bridges dissociate randomly and therefore have exponentially distributed lifetimes. Further details of the model are found in the Methods section.

**Fig. 1.**
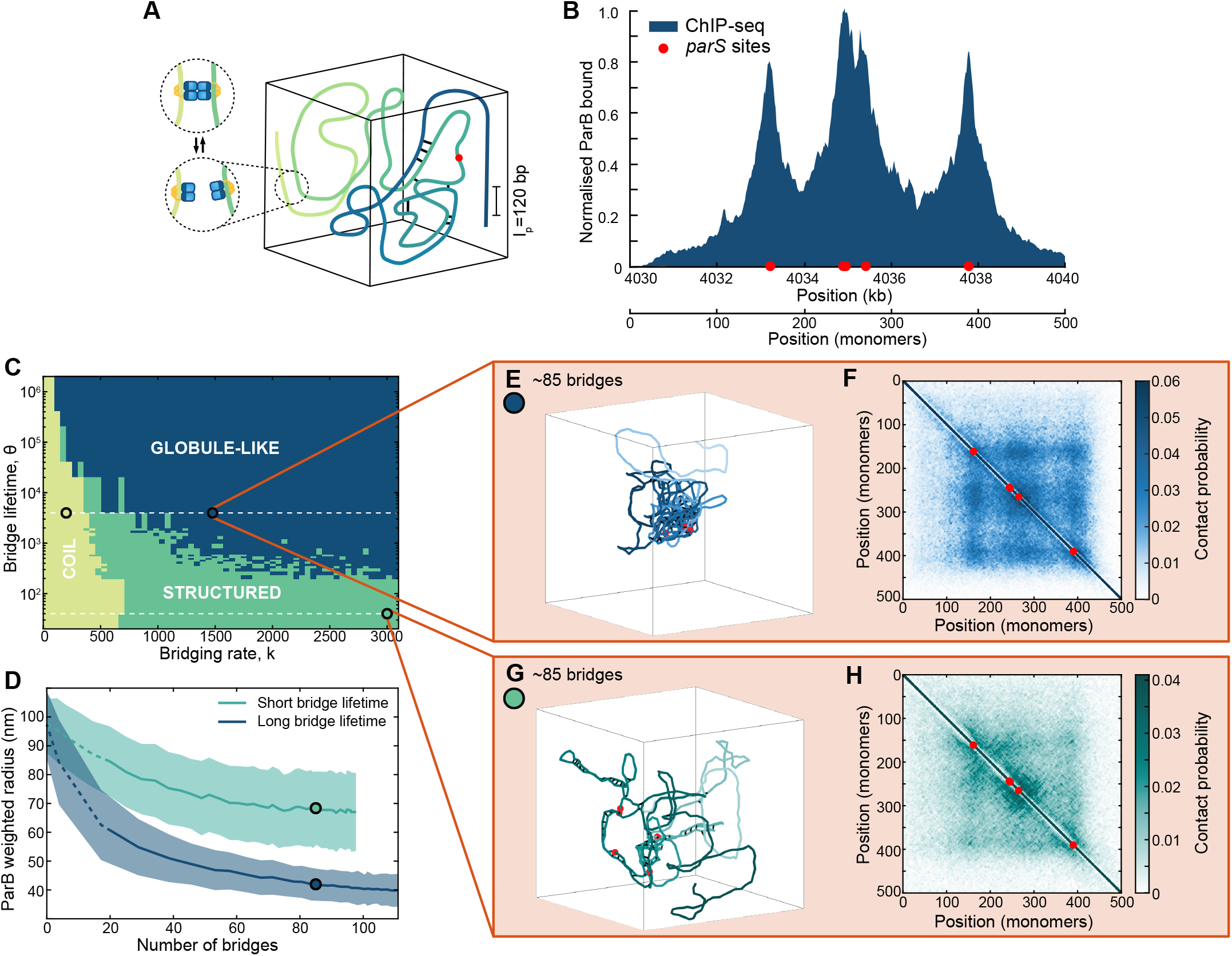
ParB bridge lifetime results in distinctly different polymer conformations. (A) Representation of polymer bridging model, showing the ParB distribution along the DNA. Bridges can form between genomically distant monomers that are in spatial proximity with a probability proportional to the occupancy of ParB at each monomer. The ParB distribution is shown explicitly in (B). (B) ParB distribution as found in (12) normalised to form a ParB bridging probability distribution. (C) Phase diagram of the system in terms of *k*, the bridging rate, and *θ*, the bridge lifetime in units of the reciprocal of the move rate (see Methods). At low bridging rates the polymer is free and in the open coil regime. With increasing numbers of ParB bridges (a threshold of 20 is used), two regimes emerge, distinguished by the size of the partition complex (the ParB weighted radius). In the globular phase the polymer forms a tight compact structure as in (E), while in the structured phase the polymer has a more open conformation defined by the formation of extended structures as in (G). (D) The ParB weighted radius for short and long bridge lifetimes (indicated by the dashed lines in (C)) as a function of the number of bridges rather than the bridging rate *k* so that the curves are comparable. Long bridge lifetimes result in a significantly lower ParB weighted radius. Shading indicates the standard deviation. Data from 1000 conformations for each parameter set. Circles indicate the respective locations of (E) and (F), and (G) and (H). (E) An example conformation of the polymer in the globular state. The location on the phase diagram is marked with a dark blue circle. (F) Average contact map at the same location based on 1000 conformations. A contact is defined as two monomers being with 5 lattice sites of one another. (G) An example conformation of the polymer in the structured state. The location on the phase diagram is marked with a green circle. (H) Average contact map at the same location, otherwise as in (F). The locations studied in (E-F) and (G-H) both have an average of ∼85 bridges. Equivalent plots for the coiled regime, indicated by the leftmost circle in (C) are shown in SI Fig. 1F,G

Since ParB dimers can slide along the DNA, the spreading of ParB throughout the centromeric region can occur, at least in principle, independently of any 3D structure. We therefore initially model ParB dimers implicitly, using the relative probability of ParB occupancy, obtained from the experimental binding (ChIP-Seq) profile (12), to specify the probability of a bridge forming when two given monomers come into proximity. This will allow us to first investigate how the observed ParB genomic distribution can, through bridging, affect the structure of the centromeric region, separately from the question of ParB spreading. We will examine the coupling between the two processes of sliding and bridging later.

### ParB bridge lifetime results in distinctly different polymer conformations

The multi-peaked ParB binding profile of *C. crescentus* consists of three clear peaks centred on five *parS* sites and covering a centromeric region of about 10 kb (Fig. 1B). While two other putative *parS* sites have been identified (12), they are not associated with significant enrichment of ParB. This profile is used in our polymer model to specify, up to an overall factor, the probability of a ParB dimer being bound to any given DNA monomer and thus specifies, again up to an overall factor, the bridging probability. Simulating the system, we found a surprising result: ParB-induced bridging leads to two distinct phases for the partition complex. Long bridge lifetimes cause the polymer to collapse into a globule-like structure (Fig. 1E), whereas at shorter bridge lifetimes the polymer is more structured with long extended localised regions of bridging (Fig. 1G). Note that ‘long’ and ‘short’ are relative to the timescale of the dynamics of the DNA polymer. Since we do not have an experimental estimate of this timescale at the lengthscale (20 bp) consider here, we cannot provide specific values.

The effect was also apparent in maps showing the location of the ParB bridges (SI Fig. 1B,D). Whereas bridge maps of the structured conformations show distinctive ±45° lines, those of the globular regime display a more random distribution. However, despite the clear differences in their conformations, both regimes exhibit very similar bridge maps at the population level (SI Fig. 1C,E), which display a checkerboard pattern centred on the *parS* sites and have no ±45° lines detectable. Such a pattern is consistent with an overall preference for contacts within and between regions associated to peaks in the ParB binding profile. A similar pattern was also observed in contact probability maps (Fig. 1F,H), though the globular regime had more contacts for the same number of bridges, as expected from its greater level of compaction. This highlights how the population-average view of DNA organisation may not be informative of the structure of individual conformations.

To better characterise these different regimes we constructed the phase diagram of the system (Fig. 1C). Three regions could be identified: a free coil regime in which there are very few bridges (less than 20) and the polymer behaviour is dictated simply by its stiffness and volume-exclusion (SI Fig. 1F,G), and the structured and globular (unstructured) regimes. We defined the transition between the structured and globular regimes using the ParB weighted radius, i.e. the radius of the *spatial* ParB distribution due to the polymer conformation (see methods). The globular state has a much smaller ParB weighted radius compared to the structured state with with the same number of ParB bridges (Fig. 1D). This radius plateaus as the system goes further into the globular regime. We therefore chose a threshold of 55 nm to distinguish the two regimes based on two standard deviations above the plateaued mean value (SI Fig. 1H).

We propose that these two condensed regimes arise due to the degree of movement the polymer makes between bridging events. Consider a polymer conformation with bridges formed between monomers in close contact. At short bridge lifetimes the polymer does not move significantly between bridging events, so that new bridges are most likely to form near to existing bridges since genomically-distant segments of the polymer are already in close proximity at these locations. Repetition of this process then leads to the extended regions of bridges which we observe. On other hand, at long bridge lifetimes the polymer re-organises between bridging events so that new close contact events between previously distant segments are more likely to occur. Therefore, the probability of bridges forming distantly from existing bridges is greater than for short bridge lifetimes. This results in a more random distribution of bridges (SI Fig. 1B) and a globular polymer configuration.

Since the globular regime is reminiscent of previous proposals for partition complex organisation (11, 20, 21), we will focus next on examining the structured state. We will return to the globular state in the final section.

### Short-lived ParB bridging leads to the formation of hairpins and helices

The structured regime found at short ParB bridge lifetimes is defined by the presence of two distinct structures, hairpins and helices. Hairpins form by the polymer bending 180° back on itself to form bridges between anti-parallel segments, whereas in helices, the polymer turns a full 360° with bridges between parallel segments (Fig. 2A). These two structures are visually different but also have different underlying bridging patterns which allows them to be clearly identified in bridge maps. Hairpins correspond to +45° lines whereas helices correspond to -45° lines. The location of the line relative to the main diagonal indicates the length of the loop of the hairpin or the period of the helix. Unsurprisingly, these structures generally form near to the *parS* sites. However, we observed substantial variation: the tip of a hairpin (indicated by where the 45° line in the bridge map intersects the main diagonal) was often reasonably far from the nearest *parS* site (Fig. 2A). At lower levels of bridging, these structures most frequently form within the region covered by the central peak containing three *parS* sites.

**Fig. 2.**
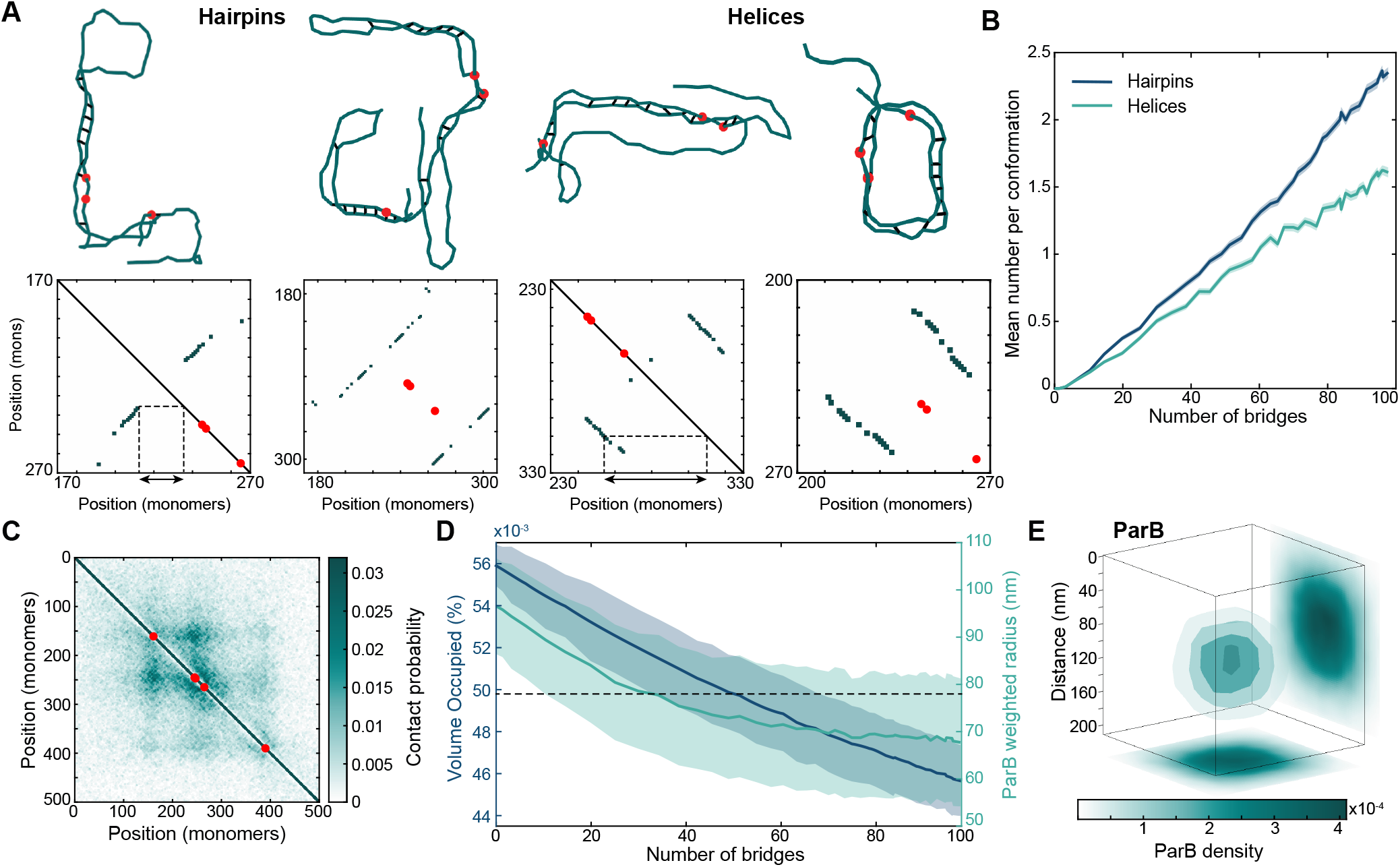
Short-lived ParB bridges results in the formation of hairpins and helices. (A) Example structures with corresponding bridge maps (bridge maps have been dilated to make lines clearer) for hairpins and helices with an average of 30 bridges. Full polymer conformations and bridge maps can be seen in SI Fig. 2A. Red dots indicate the *parS* sites. Note that as two of the sites are only separated by 42 bp they are not always distinguishable. (B) Mean number of hairpins and helices per conformation. Shading represents the SEM. (C) Average contact map for the polymer at an average of 30 short-lived bridges. (D) Volume occupied by the polymer and the ParB weighted radius. Shading represents the SD. The dotted line at 78 nm shows the experimentally determined ParB radius for *C. crescentus*. (E) Three-dimensional ParB density from partition complexes with an average of 30 bridges showing a radius of 78 nm.

We made use of the distinctive ±45 ° lines to quantify the occurrence of hairpins and helices as a function of the degree of bridging in the system (Fig. 2B). We found that the frequency of both structures increased approximately linearly with the number of bridges, with hairpins being the most common. From about ∼30 bridges every conformation contained at least one structure (SI Fig. 2B). At the highest levels of bridging studied (∼100 bridges) each conformation contained 3-4 structures, which could be of either type and involve multiple and distant *parS* sites. Nevertheless the different constituent structures could still be identified from the ±45° lines in the bridge maps. However, as discussed in the previous section, the ±45° lines are not apparent in the ensemble average contact map or bridge map which display a checkered pattern centered on the *parS* sites (Fig. 2C and SI Fig. 2C).

Consistent with *in vitro* experiments, ParB bridging led to the condensation of the DNA polymer. Both the volume occupied (see methods for definition) and the squared radius of gyration decreased with the number of bridges (Fig. 2D, SI Fig. 2D). *In vivo* the nucleoprotein complex is visualised through the spatial distribution of a fluorescently tagged variant of ParB, which forms distinct foci within cells. To connect with these observations, we combined the genomic distribution of ParB on the DNA (based on the ChIP-seq profile), with our simulated conformations of the DNA polymer to obtain the resulting spatial distribution of DNA bound ParB (Fig. 2E). The resultant spherical density was reminiscent of that observed experimentally using single molecule microscopy. The radius of the partition complex of *C. crescentus* has been measured experimentally using single molecule microscopy to be ∼78 nm (34). This could be achieved in our simulations with just 30 ParB bridges. This corresponds to a 20% decrease compared to the value in the absence of bridging (Fig. 2D).

### ParB sliding model can reproduce the multi-peaked ChIP-seq profile

In the last sections, we ignored the question of how the genomic distribution of ParB is formed but rather focused on how the observed distribution can affect, through bridging, the organisation and compaction of the centromeric region. In this section, we do the opposite and consider how ParB spreads along the DNA, while ignoring any potential effect of ParB bridging. Several recent *in vitro* studies have shown that ParB dimers of chromosomal ParABS systems (*C. crescentus, Myxococcus xanthus and Bacillus subtilis*) can entrap DNA at *parS* before sliding away in either direction in a manner akin to a DNA clamp (13, 23–27, 29). Dissociation is believed to be primarily due to CTP hydrolysis. We recently developed a stochastic model of this spreading mechanism in the context of *M. xanthus* and found that loading at *parS* sites, 1D diffusion along the DNA and dissociation was indeed able to qualitatively reproduce the observed ParB binding profile from ChIP-Seq. The predicted 1D diffusion coefficient also agreed with single molecule microscopy measurements. However, the binding profile of *M. xanthus* is relatively noisy and consists largely of a single peak centred on a cluster of all but one of its 24 *parS* sites. Therefore, the multi-peaked and less noisy profile of *C. crescentus* may serve as a better test of the *in vivo* relevance of the loading and sliding model.

We use the same fundamental model as previously (13), modified for *C. crescentus*. ParB dimers load onto the DNA, modelled as a 1D lattice, at any of 5 *parS* sites. Loaded dimers then diffuse along the lattice with effective diffusion coefficient *D* and dissociate randomly at a rate *k*_off_ (Fig. 3A). While bound they act as obstacles for other bound dimers i.e. dimers cannot move past one another. The *parS* sites must also be free for a dimer to load. The total number of ParB dimers is fixed at the measured value of 360 (34), with unbound dimers treated as a well-mixed bulk (cytosolic) population. The relative loading rate at each site is specified by its relative affinity for ParB (12), leaving a single overall loading rate *k*_on_.

**Fig. 3.**
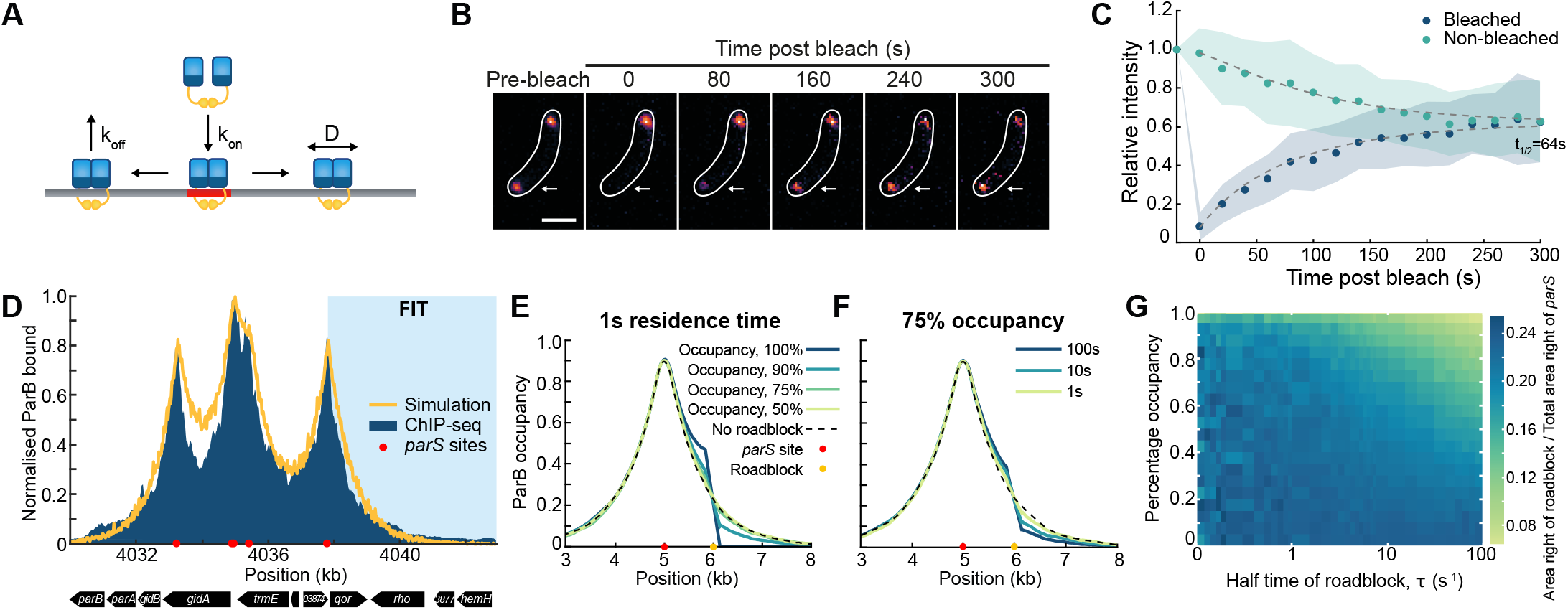
ParB sliding can reproduce the multi-peaked *C. crescentus* profile. (A) Diagram representing the sliding model, showing ParB dimers binding at a *parS* site, diffusing along the lattice and unbinding. (B) Representative images of fluorescence recovery after photobleaching (FRAP) experiment. A single eGFP-ParB focus (arrow) was photobleached in a cell containing two foci. All timelapse images for this cell can be seen in SI Fig. 3A. (C) Analysis of FRAP data. The average relative intensity of the bleached (blue) and unbleached (green) partition complex (±SD) are shown as a function of time (n=41 cells). Dashed lines represent the behaviour from a fitted model, which found a half-time of 64 s (see methods). (D) Simulation of ParB sliding compared to ChIP-seq data from (12), both normalised by maximum height. Shaded area indicates the part of the ChIP-seq profile that was fitted to an exponential to find the effective diffusion coefficient. (E) Simulations of ParB sliding at 1s residence time for different roadblock occupancies. The roadblock is indicated by the yellow dot. Similar figures for 10 and 100s residence times are in SI Fig. 3D. (F) Same as in (E) but for 75% occupancy with a varying roadblock lifetime. In both (E) and (F) the dotted line shows the profile when there is no roadblock. (G) Phase diagram displaying the difference between simulations with and without a roadblock. The colour indicates the ratio of the area of the distribution to the right of the roadblock to the total area on the right hand side of the *parS* site. The locations of the roadblock and the *parS* site are shown in (E) or (F).

We first determine the effective diffusion coefficient *D* and the dissociation rate *k*_off_. To estimate the latter, we performed fluorescence recovery after photobleaching (FRAP) of eGFP-ParB foci in pre-divisional cells containing two foci (partition complexes) (Fig. 3B). After bleaching one of the two foci, the fluorescence signal recovered with a half-time of 64 s (Fig. 3C, SI Fig. 3A,B). This provides an estimate for the dissociation rate *k*_off_ (see methods). To determine the diffusion coefficient, we fit the outer part of the third peak to an exponential *e*^*x*/*λ*^ with 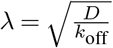 (Fig. 3D), the predicted continuum distribution under this model for an isolated *parS* site (see methods). The fitted value of *λ* = 710 bp, then gives *D* = 5600 bp^2^s^−1^ = 610 nm^2^s^−1^.

One model parameter remains to be determined - the overall loading rate of ParB *k*_on_. Previous measurements have estimated that approximately 80% (290) of ParB dimers in the cell are in ParB foci. In contrast, we find that even at high loading rates less than 220 ParB dimers are associated with the DNA (SI Fig. 3C). Increasing *k*_on_ further does not substantially increase the number of ParB bound as the *parS* sites are almost continuously occupied. The disparity in the number of DNA-associated dimers may be due to several factors. Firstly, the maximum possible number of associated dimers in our simulations is dependent on the chosen discretisation since each lattice site/monomer can be bound by a single ParB dimer. Thus if the footprint of ParB is smaller than our discretisation size of 20 bp, we would be underestimating the achievable ParB occupancy. Secondly the *in vivo* estimate of the cellular ParB concentration is based on quantitative Western blotting, which has a substantial margin of error (38). ParB foci may also contain a cytosolic or non-specifically bound population that is not accounted for in our model.

Given the above, we choose the loading rate for our model by finding the best fit of the simulations to the ChIP-seq data (SI Fig. 3C), obtaining *k*_on_ = 200 · *k*_off_. This results in remarkably good agreement between the model and the ChIP-Seq profile (Fig. 3D), indicating that loading and diffusive sliding of ParB dimers can indeed explain the observed binding profile. It also suggests that dimers are largely unaffected by transcription and other processes that could hinder ParB spreading since we have not accounted for these effects in our model. This may not be the case for other systems such as F plasmid that show changes in the binding profile coincident with promoters (21). Indeed, *in vitro* experiments have shown that high-affinity DNA-binding proteins, such as EcoRI (with the catalytically-inactive E11Q mutation) and TetR, can block the sliding of ParB dimers along the DNA (24, 26, 29).

### Residence time and percentage occupancy of road-blocks impacts their effect on ParB sliding

To better understand how roadblocks can impact the spreading of ParB dimers, we used our sliding simulation to examine the effect on spreading from a single *parS* site. Representative of the biological situation, we do not consider a permanent road-block but rather a dynamic one, which we specify in terms of its average lifetime and occupancy (the fraction of time the roadblock is present). We found that at a lifetime of 1 s, the roadblock had a surprisingly mild effect on spreading, only becoming noticeable from an occupancy of about 75%. Even at 95% occupancy, roughly half the number of dimers slide past the site of the roadblock as in its absence (Fig. 3E). Similarly, at 75% occupancy, a negative effect on spreading was only observed for roadblock lifetimes greater than about 1 s (Fig. 3F).

We can understand these results as follows. When the road-block is present for a time much shorter than the time interval between dimer crossing attempts then a backlog of dimers does not develop. Even for longer times, the backlog of dimers can be cleared if there is enough time between roadblock events i.e. if the average roadblock occupancy is sufficiently low (Fig. 3G). These results may explain why we observe no significant deviation of the ParB binding profile from that expected from our simple loading and sliding model - the *in vivo* occupancy and residence times of proteins binding to the centromeric region of *C. crescentus* may simply not be large enough to substantially affect ParB spreading.

### Coupled simulations of sliding and bridging

We next investigate whether ParB bridging is compatible with the ParB binding profile i.e. would the spontaneous formation of ParB bridges between spatially-proximal but genomically distant ParB dimers limit overall ParB spreading and produce a fundamentally different binding profile? To answer this question we coupled our polymer and sliding simulations together (Fig. 4A). We assume that bridged dimers are not able to slide along the DNA, due to the entrapment of genomically distant regions, so that they act as roadblocks for unbridged sliding dimers. The procedure is as follows: The location of each ParB dimer from the sliding simulation is given as an input to the polymer simulation, which is then ran until the next bridging or unbridging event occurs (at some time Δ*t* later). The updated configuration is then returned to the sliding simulation, which is subsequently ran for the same time Δ*t* and the cycle repeats. The simulations are ran to equilibrium and the ParB distributions and polymer conformations are recorded.

**Fig. 4.**
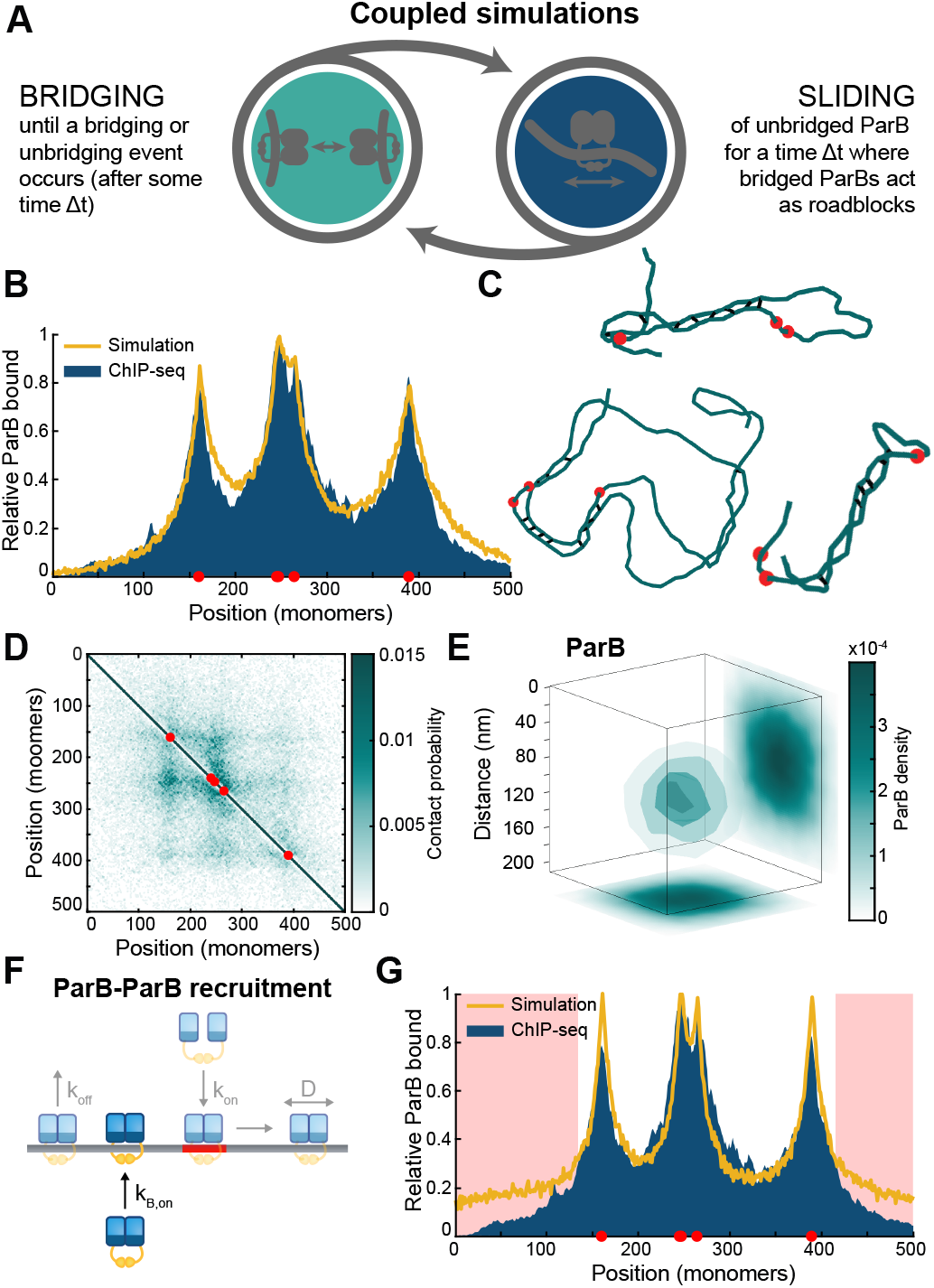
Sliding is not inhibited by short-lived ParB bridges. (A) Representation of coupled polymer simulations in which we combine bridging and sliding. (B) Profile of ParB as generated from sliding and bridging simulations with a bridge lifetime of 1s compared to ChIP-seq data from a previous study (12), both normalised by maximum height. (C) Examples of hairpin and helical structures found in coupled simulations, with an average of 25 bridges. Full conformations and individual bridge maps can be found in SI Fig. 4A. (D) Average contact map for coupled simulations (top right) and uncoupled simulations (bottom left) with a radius of ∼78 nm. Each contact map is made from 1000 simulations. (E) Three-dimensional average of the ParB partition complex for 25 bridges with a radius of 78 nm. (F) Diagram displaying the addition of ParB-ParB *in cis* recruitment to the coupled model. (G) Profile of ParB along the polymer with additional ParB-ParB recruitment, areas where the profile is substantially different to ChIP-seq data are highlighted in red. For all plots shown the simulations are in the structured regime.

The same values determined in the previous section are used for the ParB dimer loading and dissociation rate. There are currently no estimates for the bridge lifetime. We expect bridges to have a significantly shorter lifetime than that of ParB dimers on the DNA and therefore a nominal value of 1 s is chosen. With too high a value (of the order of the ParB lifetime on the DNA) sliding ParB dimers would not have time to move past roadblocks (ParB bridges) before unbinding. This leaves the sliding diffusion coefficient as the free parameter. We are unable to use the value found for the diffusion coefficient in the previous section due to the introduction of ParB bridges resulting in ParB dimers sliding a shorter distance creating sharper peaks. Instead we choose the sliding diffusion coefficient such that we recover the expected genomic distribution. Thus this value must be tuned based on the number of ParB-ParB bridges.

With the bridge lifetime fixed we access the two regimes discussed in the first section through the mobility of the polymer. First, for the structured regime, we found that the coupled simulations could reproduce the binding profile measured by ChIP-seq (Fig. 4B), with an even better fit than we obtained from the non-polymeric sliding simulation (Fig. 3D). Importantly, we also observed the same hairpin and helical structures as in the uncoupled polymer simulations that had the ParB binding profile given as a input (Fig. 4C) and obtained very similar average contact (Fig. 4D) and bridge maps (SI Fig. 4B). These structures again compact the polymer and we could achieve the measured radius of 78 nm (Fig. 4E).

We could also reproduce the ChIP-seq profile in the globular regime (SI Fig. 4C) with similar contact and bridge maps (SI Fig. 4D and E). Interestingly, the ParB weighted radius at the same number of bridges was significantly larger in the coupled simulations than in the uncoupled simulations. This suggests that sliding of ParB pushes the system towards the structured regime (SI Fig. 4F and G).

Recent *in vitro* studies have shown that DNA-loaded ParB dimers of *B. subtilis* can load additional dimers independently of *parS* (‘ParB-ParB recruitment’) (39), potentially explaining the cooperative non-specific DNA binding observed previously (15, 18, 19) and consistent with interactions between dimers through their N-terminal domains (16, 40). To explore whether such recruitment could be relevant *in vivo*, we added *in cis* ParB-ParB recruitment to our model (Fig. 4F). Although *in trans* recruitment was also shown by the same authors this would be substantially more challenging and computationally intensive to implement.

We found that even a relatively low ParB-ParB recruitment rate, for which the total number of bound dimers increased by less than 20%, resulted in a fundamentally different binding profile. ParB spreading was increased through the appearance of slowly decaying ‘shoulders’ at the extremes of the distribution. As a result the distinctive exponential decay seen in the experimental ChIP-seq profile is no longer reproduced (Fig. 4G). As suggested in (39) where they see ParB-ParB recruitment accounting for only 10% of ParB binding events, our results suggest that ParB-ParB recruitment does not play a significant role *in vivo* in ParB spreading.

## Discussion

The sliding and bridging model presented here uses recent discoveries to probe the formation and structure of the partition complex. Recent *in vitro* based studies have shown that, dependent on CTP, ParB can load onto DNA at *parS* sites before sliding randomly along the strand (13, 23–27). It was also shown that ParB can efficiently condense DNA, again in the presence of CTP, through the formation of bridges between genomically distant DNA regions (10, 15, 28, 29). While we have not explicitly modelled the CTP dependent nature of these processes, our model is consistent with CTP hydrolysis triggering the unbinding of ParB dimers and therefore setting the length scale of sliding (13, 24). Our model predicts that the dynamic sliding and bridging of ParB results in two different conformational regimes, one globular, one structured for long and short bridge lifetimes respectively. The latter regime is dependent on the stiffness of the DNA. If that is ignored, short range bridges between next to neighbouring monomers dominate and DNA structures do not develop. We showed how the genomic distribution of ParB could define its spatial distribution through the formation of ParB bridges. We then explicitly modelled both the sliding of ParB along the DNA and the formation ParB-ParB DNA bridges. Importantly, we found that sufficiently short-lived bridges do not hinder sliding of ParB along the DNA and our model could reproduce both the measured genomic and spatial distribution of *C. crescentus* ParB.

We speculate that the two different regimes could have relevance in different biological contexts. The hairpins and helices of the structured regime may facilitate the loading of SMC (structural maintenance of chromosomes) complexes onto the DNA (41). While this is known to be due to ParB at the *parS* sites (42), the precise mechanism is a topic of ongoing study (43, 44). However, ParB mutations that eliminate SMC recruitment are also known to reduce the ability of ParB to form a higher-order nucleoprotein complex (43, 45, 46). This leads us to postulate that the ParB-induced DNA structures we observe are relevant for the loading of SMC complexes (Fig. 5). Indeed, in the absence of a specific mechanism, DNA hairpins are unlikely to form given the intrinsic stiffness of the DNA. Furthermore, chromosomal ParAB*S* systems often have multiple separated *parS* sites (6) that produce a multi-peaked binding profile (9, 10, 12, 13, 30–33), whereas a single cluster of *parS* sites appears to be more common for plasmid-based systems (47). Separated *parS* sites would allow the formation of multiple hairpins and could thereby be beneficial for SMC loading.

**Fig. 5.**
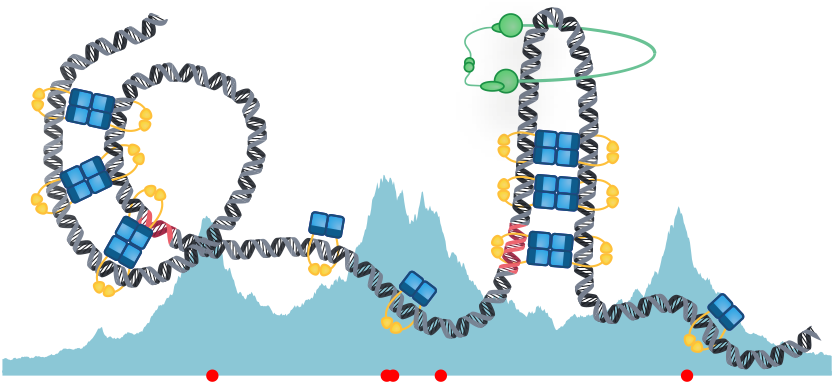
A sliding and bridging model can reproduce the genomic and spatial distribution of ParB, forming hairpins and helical structures which organise the DNA.

In contrast, ParABS-carrying plasmids, especially those of *E. coli* and other bacteria that do not carry SMC (48), would likely not require these structures. Instead it may be advantageous to form a more compact partition complex to better facilitate the partitioning function of ParABS. Indeed, while F plasmid ParB spreads over a four times larger region than ParB of *C. crescentus* (11), the resultant partition complex is significantly smaller (a radius (2*σ*) of 35 nm) (49). Thus, we speculate that plasmid-based ParABS systems may operate in the more compact globular region. Super-coiling, which is not accounted for in our model, may also contribute (22). The conformations we observe in our simulations are similar to those recently seen using atomic force microscopy for *B. subtilis* ParB (29) but more detailed study is required. Our model could also be better characterised by knowledge of the ParB-ParB bridge lifetime, which could be achieved *in vitro* by using magnetic tweezers to probe the relaxation time of ParB-condensed DNA upon removal of ParB from the buffer. *In vivo* characterisation of the partition complex is more challenging. While our simulated contact maps are in principle comparable to the experimental contact maps produced by chromatin conformation capture (HiC), the resolution of this technique is not yet sufficient to probe DNA structure at the short lengthscale of the *C. crescentus* centromeric region. This may change as the technique improves (50, 51).

Overall, we have presented a physical model for the formation of the partition complex of the ParABS system. Our dynamical sliding and bridging model reconciles the recent result that ParB spreads along the DNA by sliding like a DNA clamp with the ability of ParB to condense the centromeric region into a nucleoprotein complex. Future experimental work will help in evaluating the model and testing its predictions.

## Methods

### A. Polymer model

We simulate the DNA of the centromeric region using the Bond Fluctuation Model (BFM) (36). Specifically, the DNA is modelled as a linear chain on a 3D cubic lattice with reflective boundary conditions. Each monomer occupies a cubic site of the lattice including the eight associated lattice points. Furthermore, each lattice point can only be occupied by one monomer at a time to account for the excluded volume of the chain. Individual monomers are connected by bond vectors taken from a set of 108 allowed vectors. This set is chosen such that the polymer chain cannot pass through itself (36). Monomers can move one lattice site at a time in each Cartesian direction subject to the constraint on allowed bond vectors and the excluded volume. The model is ergodic in that the configuration space of the polymer can be sampled using only local moves. We use a kinetic implementation, based on the Gillespie method (52), as this allows us to incorporate the dynamics of bridging (see below).

We take each monomer to represent 20 bp since this is the approximate footprint of a ParB dimer and leads to computationally tractable simulations. In *C. crescentus*, ParB spreads over a ∼10 kb region of the chromosome and we therefore simulate a polymer with a corresponding length of 500 monomers.

In order to account for the stiffness of the DNA, we follow the approach of Zhang et al. (53) and introduce a squared-cosine bending potential *E* between successive bonds

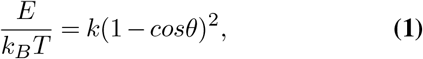

where *θ* is the change in angle between successive bonds and *k* is a parameter controlling the stiffness. When a monomer attempts a move, this leads to a change in three bond angles: the angle at the monomer and at its two neighbours. An attempted move is then accepted with probability *P* = *min*(1, exp(−Δ*E/k*_*B*_*T*)), where Δ*E/k*_*B*_*T* is the change in energy due to the move. We set the stiffness parameter *k* = 14 as this gave a persistence length (calculated according to the angle between consecutive bonds (53)) of 120 bp (SI Fig. 1A) inline with recent experimental measurements (37).

Similar to other bacteria, chromosomal DNA in *C. crescentus* constitutes a volume fraction of about 1-2%. We obtain this volume ratio in the simulations by setting the size of lattice appropriately. In the BFM the volume occupied by the polymer is not a fixed quantity due to the large set of bond vectors - the excluded volume associated to each monomer can overlap. However, we can measure the occupied volume by dilating the three-dimensional binary image describing the occupancy of each lattice site using a cubic structuring element of width 3 (we use the MATLAB function *strel*). This gives precisely the excluded volume of the entire polymer (recall that each monomer uniquely occupies eight lattice points). Using a 90×90×90 cubic lattice and the stiffness parameter chosen above, we find an excluded volume fraction of 1.7% (an excluded volume per monomer of ∼22 lattice sites).

Upon this stiff polymer framework, we implement bridging between non-neighbouring monomers. Our implementation is similar to that of Bohn and Heermann (54). Any two non-neighbouring monomers that are within (strictly less than) a spatial distance of 3 lattice sites are allowed to bridge at a given rate. The rate (probability) of bridging depends on the positions of the monomers within the polymer. In Figures 1 and 2, the rate is specified, up to an overall factor, by the ParB binding profile (determined by binning the experimental ChIP-Seq profile at the 20 bp resolution of the polymer). The rate of bridge formation between two monomers that are in proximity is then proportional to the product of the ParB occupancy at each site. Bridged monomers can still move on the lattice but must maintain a bridge length strictly less than 3. Each monomer can only bridge with at most one other monomer.

Bridge lifetimes *λ* are exponentially distributed with a mean of 1 s. Acceptable monomer moves (moves that obey the volume exclusion, bond length, bridge length and stiffness conditions) are attempted with a rate *p*. When discussing the bridge lifetime in terms of *θ* we refer to the number of moves made during the bridge lifetime, *θ* = *pλ*. We vary *θ* via the move rate *p*. For the phase diagram presented in Fig. 1C we vary *p* from 20 to 2 × 10^6^, for the rest of the paper we take the arbitrarily chosen values of *p* = 40 to represent the structured regime and *p* = 4 × 10^3^ for the globular regime, shown by the white dotted lines in Fig. 1C.

For any given parameter set, simulations are first run until equilibrium is reached as determined by the volume occupied by the polymer reaching an approximate constant value. We use the volume occupied rather than the usual squared radius of gyration as the former was found to be a much less noisy measure. The conformation of the polymer is then recorded. We repeat this process for 1000 random initial configurations.

#### Calculating ParB radius

The ParB radius is calculated by combining the genomic distribution of ParB on the DNA (either based on the ChIP-seq profile for the uncoupled polymer simulations or the simulated position of ParB dimers in the coupled simulations) with the simulated conformations of the DNA polymer to obtain a spatial distribution of DNA bound ParB. We take an average across all 1000 conformations, aligning them by their centroids, to obtain a 3D density. We then determine the radius within which 95% of ParB dimers are found. We convert this value from lattice units to nanometers as follows. In our (stiff) polymer simulations, the bond length between monomers varies but has an average value of 3.0 lattice units. Since every bond/monomer corresponds to 20 bp and the length of a base pair is 0.33 nm (55), a lattice unit corresponds to 2.2 nm.

### B. Model of ParB sliding

We model the loading, sliding (diffusion) on and unbinding of ParB dimers from the DNA using the same approach as in our recent work on *M. xanthus* (13). The DNA is modelled as a one-dimensional lattice with each lattice site corresponding to 20 bp. The model is single-occupancy - loading and sliding can only occur if the target lattice site is free. ParB dimers can load at some number of special lattice sites, corresponding to the *parS* sites. For the simulations of sliding in *C. crescentus*, the relative loading rate at each *parS* site is determined by 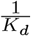 here *K*_*d*_ is the measured dissociation constant (12). The loading rate at each site is then determined by multiplying by an overall factor *k*_on_. Dimers diffuse to unoccupied neighbouring lattice site with a rate *d* = *D/h*^2^, where *D* is the effective diffusion coefficient and *h* is the lattice spacing. Unbinding occurs randomly with rate *k*_off_. The total number of dimers is fixed as 360 as estimated for *C. crescentus* (34). Any unbound dimers are assumed to be in the cytoplasm which we take to be well-mixed.

For each parameter set the simulation is first run until steady state is reached and then the distribution of ParB is recorded at regular time intervals, sufficiently separated to be independent samples of the equilibrium distribution.

#### Analytical description

We provide an analytical description of sliding for the simplified case of a single *parS* site. Consider ParB dimers diffusing on an infinite single-occupancy lattice (lattice spacing *h*). Dimers can move to any unoccupied neighbouring site at a rate *d*. Dimers load onto the lattice at a site *i* = 0 with rate 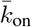 and unbind with rate *k*_off_. We denote the probability of there being *n* ParB dimers at the *i*-th site by *P*_*n*_(*i, t*) (*n* = 0, 1 due to single occupancy). The chemical master equation which corresponds to this system of reactions is

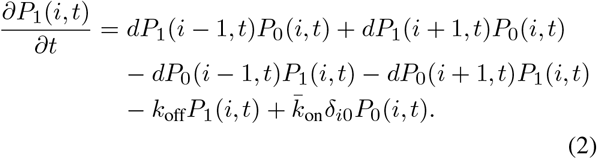

Using *P*_0_(*i, t*) + *P*_1_(*i, t*) = 1, we can rewrite this in terms of the expected number of dimers at each site, 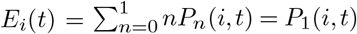, as

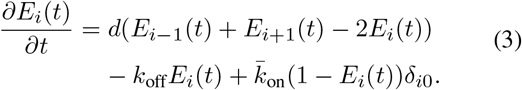

A similar equation for *E*_*i*_(*t*) is obtained for a multioccupancy model but with a different pre-factor in the Kro-necker delta term (56) i.e. the equilibrium distribution of both models have the same form. This is most easily described in the continuum limit (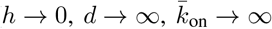 keeping *D* = *dh*^2^ and 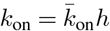 fixed) in which we obtain

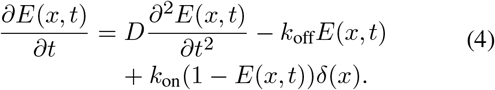

The steady-state solution of this equation is

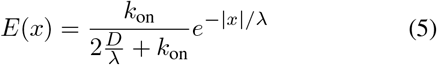

where 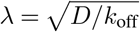 is the associated diffusive length-scale.

#### Fit to ChIP-Seq profile

Before fitting to the experimental ChIP-seq profile we first binned the profile at 20 bp resolution to match the simulations. We then fit to the right side of the right most peak of the experimental profile (Fig. 3C) to an exponential *y* = *ae*^*x/λ*^ as expected from the analysis above. We use the MATLAB *fit* function to fit for the length scale parameter *λ*, for which we find *λ* = 710 bp. The analysis above shows that 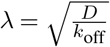 ically. We can therefore use the estimate for *k*_off_ obtained from the FRAP experiment to obtain *D* = 5600 bp^2^s^−1^ = 6.1 × 10^−4^ μm^2^s^−1^. Note that this value of *D* does not account for any roadblocks beyond the effect of sliding ParB dimers on each other. The remaining parameter of the sliding model is the overall factor of the loading rates, *k*_on_. This is determined by finding the best fit of the stochastic model to the entire ParB binding profile as determined by the mean square error between the ChIP-Seq profile and the steady-state profile obtained from the simulations (Fig. S3B). Both profiles are normalised to the same area under the curve before the mean square error is calculated.

#### Roadblock simulations

For the roadblock simulations we used the same framework but with a single *parS* site and choose a high loading rate *k*_on_ = 100 such that this site is occupied the majority of the time. We use the values of *D* and *k*_off_ as above. A roadblock is implemented as another particle that can bind and unbind to and from a specific site 25 lattice sites (500 bp) away from the *parS* site.

To explore the effect of the roadblock we either 1) fix the unbinding rate *k*_R,off_ such that the residence half-time (*τ* = log(2)*/k*_R,off_) of the roadblock remains constant and then vary the binding rate *k*_R,on_ to vary the occupancy of the road-block, or 2) fix the occupancy (*k*_R,on_*/*(*k*_R,on_ + *k*_R,off_)) of the roadblock and vary both *k*_R,on_ and *k*_R,off_ by the same factor.

### C. Coupled bridging and sliding model

In the coupled simulations the range over which the sliding simulations take place is reduced such that it is 500 lattice sites longer than then polymer where 250 lattice sites are added to either side, this is sufficient length that ParB dimers are unlikely to fall off the edges. Secondly, the bridging probability is now given by the exact ParB locations and is therefore either one or zero at any given location depending on whether a ParB is present or not respectively.

To combine the sliding and polymer simulations the location of each ParB dimer from the sliding simulation is given dynamically as an input to the polymer simulation. The polymer simulation is then run until the next bridging or unbridging event occurs (at some time Δ*t*). The updated locations of the bridged and unbridged ParB dimers are then returned to the sliding simulation, which is ran for some time Δ*t*. This cycle is repeated until a predetermined time is reached. Bridged dimers are treated as being unable to diffuse along the DNA due to the topological constraints being bound to distal DNA regions. Therefore, bridged dimers act as roadblocks for the unbridged dimers preventing them from sliding past.

We first run the sliding simulation to equilibrium, then give this as an input to the polymer simulation before allowing this to run to equilibrium. All the ParB dimers bridged at the end of this equilibrium are then updated in the sliding simulation. The simulations are then run in a coupled manner as described above, this is again run until equilibrium has been reached before a snapshot is taken. When referring to the structured and globular regimes the same nominal values, 40 and *p* = 4 × 10^3^ respectively, are used taken for *p*. For each parameter set tested 1000 different simulations are run where the starting polymer conformation and ParB distribution is different in each simulation.

### D. Fluorescence Recovery After Photo-bleaching (FRAP)

*C. crescentus* strain MT174 (parB::egfp-parB) (57) was grown in M2G minimal medium (58) at 28 °C and 220 rpm for 36 h to an OD600 of ∼0.6. Cells were spotted on pads made of 1% (w/v) agarose in M2G medium. Images were taken with a Zeiss Axio Imager.Z1 microscope equipped with a Zeiss Plan Apochromat 100x/1.46 Oil DIC objective and a pco.edge 4.2 sCMOS camera (PCO). An X-Cite 120PC metal halide light source (EXFO, Canada) and an ET-EGFP filter cube (Chroma, USA) were used for fluorescence detection. FRAP analysis was conducted by bleaching single EGFP-ParB foci using a 488 nm-solid state laser and a 2D-VisiFRAP multi-point FRAP module (Visitron Systems, Germany), with 2-ms pulses/pixel at 20% laser power. After acquisition of a prebleach image and application of a laser pulse, 16 images were recorded at 20 s intervals with VisiView 4.0.0.14 (Visitron Systems). For each time point, the average fluorescence intensities of equally sized regions containing the bleached focus, the non-bleached focus, the cell background and a reference region of the agarose pad were determined, using Fiji 1.49 (59). After background correction, normalisation and averaging of the focus intensities, the recovery half-time was calculated by fitting the data as described below.

#### D.1. Analysis

To calculate the residence time of ParB dimers from the photo-bleaching we perform a simple manipulation of the data. Following the standard calculation used in (49) the FRAP experiments can be described by a simple kinetic model for the ParB proteins in the partition complex and the ParB in the rest of the cytoplasm. Considering *B*_1_(*t*) and *B*_2_(*t*) as the average number of ParB proteins in each partition complex after photo-bleaching, *B*_tot_ the total number of ParB dimers, and *k*_in_ and *k*_out_ the rate to enter and exit the partiton complexes respectively, the system can be written as:

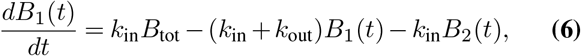

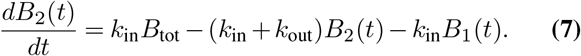

In order to fit to the data more easily we consider the sum and difference, *B*_±_ = *B*_1_(*t*) ± *B*_2_(*t*):

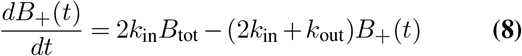

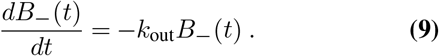

The general solution to these equations is given by:

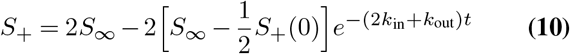

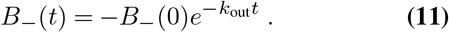

A simple exponential fit of our data to the difference curve (SI Fig. 3B) finds *k*_out_ =0.011 s^−1^, or a half time in the focus of 64 s and *B* (0) =0.91. Then fitting to the sum, taking *B* to be equal to 0.62, we find *B*_ℝ_(0) = 1.06 and *k*_in_ =0.0035 s^−1^. Using these fitted values we can plot *B*_1_(*t*) and *B*_2_(*t*) in Fig. 3C using the simple transformation 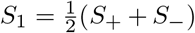 and 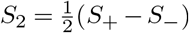.

---

**SI Fig. 1.**
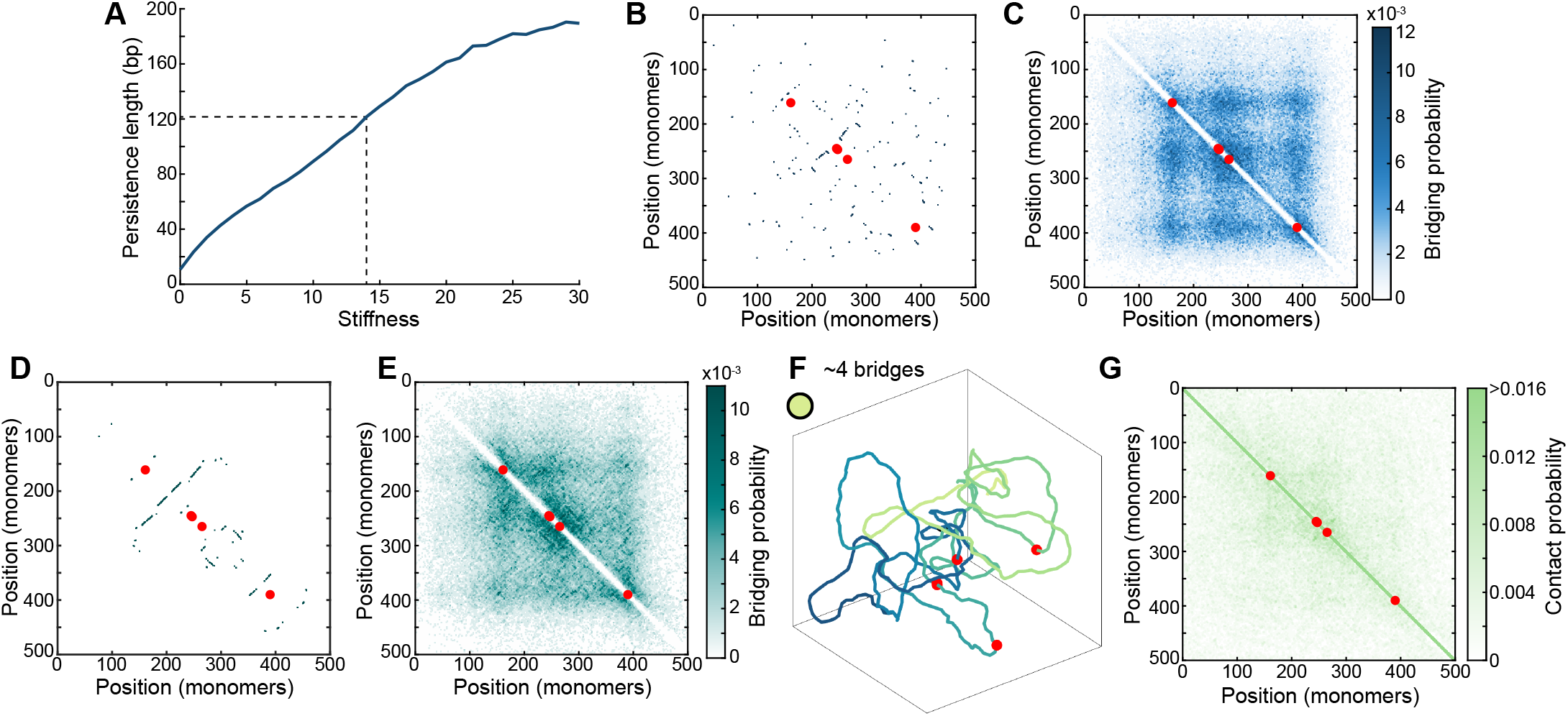
ParB bridge lifetime results in distinctly different polymer conformations. (A) The stiffness of the polymer versus the persistence length calculated. We choose a stiffness of 14 in order to give a persistence length of ∼120 bp. (B) Individual bridge map for the globular polymer conformation shown in Fig. 1E. (C) Average bridge map for the polymer in the globular regime. (D) Individual bridge map for the structured polymer conformation shown in Fig. 1G. (E) Average bridge map for the polymer in the structured regime. (F) An example conformation of the polymer in the free coil state. The location on the phase diagram in Fig. 1C is marked with a light green dot. (G) Average contact map at the same location. Based on 1000 conformations.

**SI Fig. 2.**
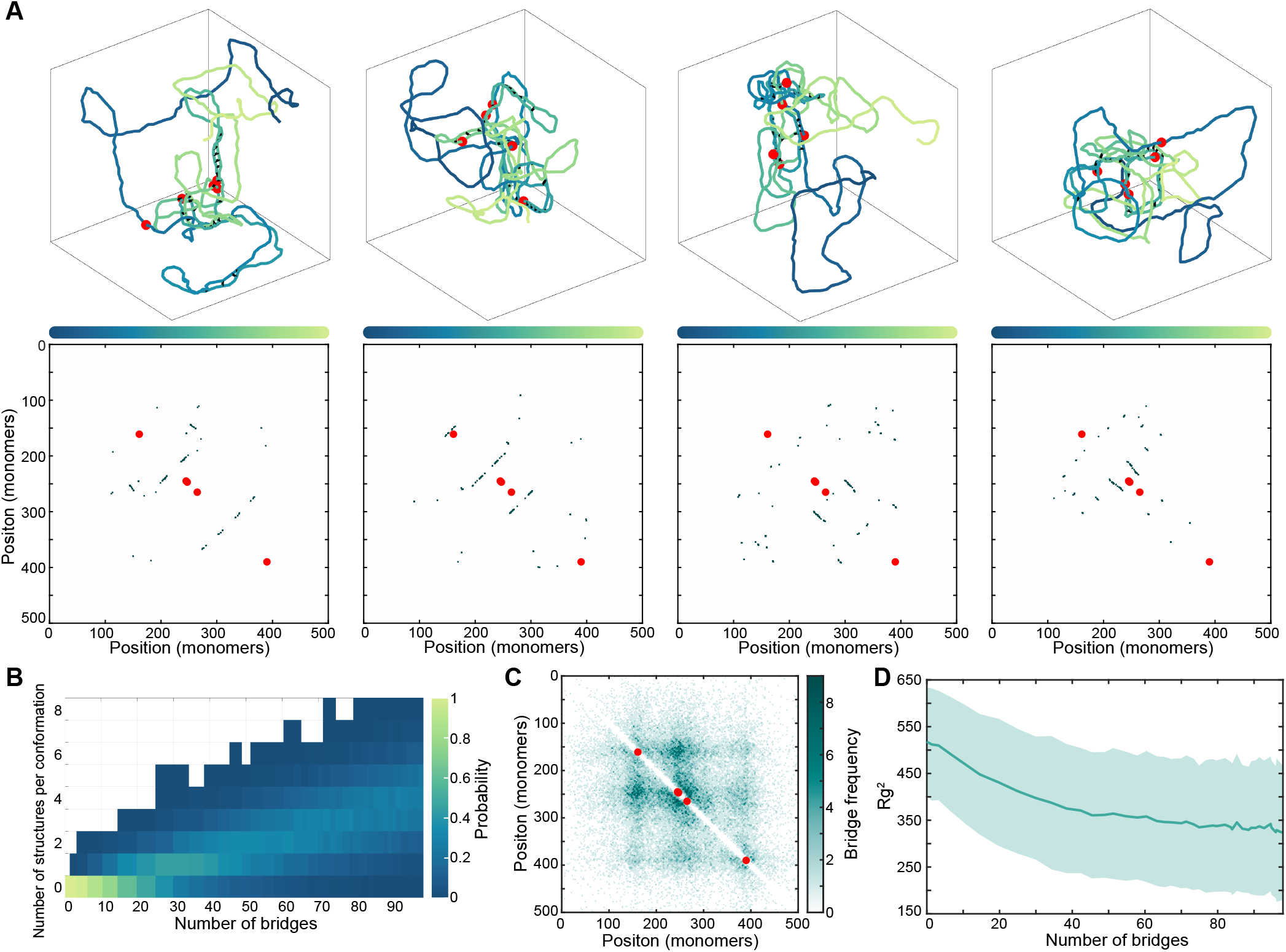
Short-lived ParB bridges result in the formation of hairpins and helices. (A) Complete polymer conformations and bridge maps for the structures shown in Fig. 2A. (B) Histogram showing the probability of a certain number of structures being present in any given conformation. After ∼ 30 bridges it becomes most likely that a given conformation will have at least one structure. (C) Bridge map for the polymer at a population level with an average of 30 short-lived bridges. (D) Squared radius of gyration (± SD) for the polymer as the number of bridges increases.

**SI Fig. 3.**
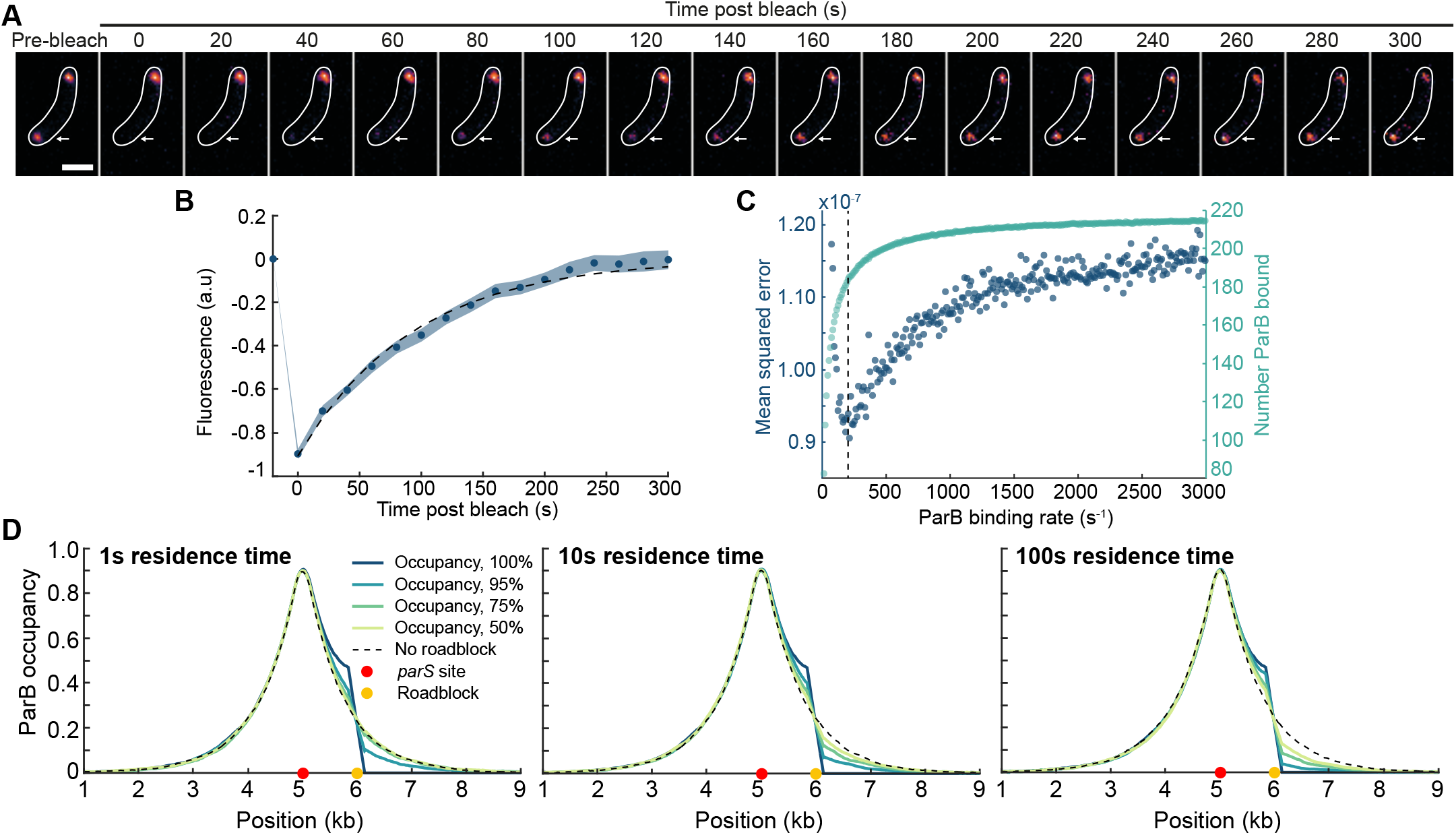
ParB sliding can reproduce the multi-peaked *C. crescentus* profile. (A) Representative images of a fluorescence recovery after photobleaching experiment. A single eGFP-ParB partition complex (arrow) was photobleached in a cell containing two condensates. (B) Difference curve *B*_−_(*t*) = *B*_1_(*t*) − *B*_2_(*t*) (blue) as described in FRAP analysis methods with corresponding fitted curve (grey) finding a half life of 64 s for ParB dimers. (C) Mean squared error for the result of simulations compared to ChIP-seq data and the number of ParB bound as the ParB binding rate is varied. Grey line marks the selected ParB binding rate, *k*_on_ =200 s^−1^. (D) Sliding profile from simulations for ParB sliding from a single *parS* site (red dot) with a roadblock located at the yellow dot where the percentage occupancy of the roadblock is varied. The residence time is varied as 1s, 10s, and 100s and displayed in each case.

**SI Fig. 4.**
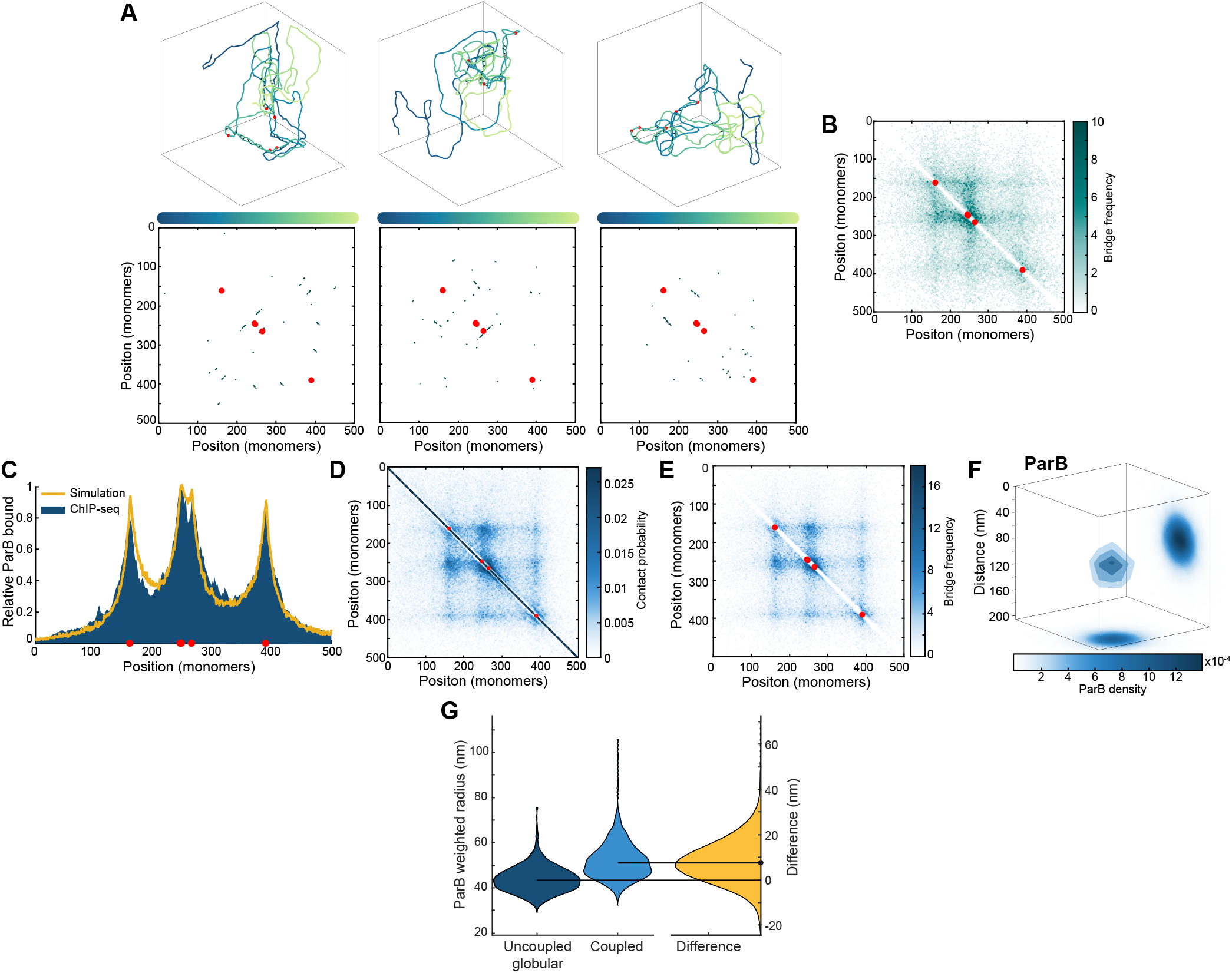
ParB sliding is not inhibited by short-lived ParB bridges. (A) Complete conformations and individual bridge maps for hairpin and helical structures found in coupled simulations in the structured regime with an average of 25 bridges. (B) Average bridge map from coupled simulations in the structured regime with an average of 25 bridges. (C) Profile of ParB as generated from the sliding and bridging simulations in the globular regime compared to the ChIP-seq data from a previous study (34). (D) Average contact map for coupled simulations in the globular regime with an average of 73 bridges. (E) Average bridge map from coupled simulations in the globular regime with an average of 73 bridges. (F) Three-dimensional average of the ParB partition complex in the globular state. (G) Violin difference plots for the ParB weighted radius in uncoupled and coupled (with explicit ParB dimers) simulations where there is an average of 73 bridges.

